# Estimating RNA structure chemical probing reactivities from reverse transcriptase stops and mutations

**DOI:** 10.1101/292532

**Authors:** Angela M Yu, Molly E. Evans, Julius B. Lucks

## Abstract

Chemical probing experiments interrogate RNA structures by creating covalent adducts on RNA molecules in structure-dependent patterns. Adduct positions are then detected through conversion of the modified RNAs into complementary DNA (cDNA) by reverse transcription (RT) as either stops (RT-stops) or mutations (RT-mutations). Statistical analysis of the frequencies of RT-stops and RT-mutations can then be used to estimate a measure of chemical probing reactivity at each nucleotide of an RNA, which reveals properties of the underlying RNA structure. Inspired by recent work that showed that different reverse transcriptase enzymes show distinct biases for detecting adducts as either RT-stops or RT-mutations, here we use a statistical modeling framework to derive an equation for chemical probing reactivity using experimental signatures from both RT-stops and RT-mutations within a single experiment. The resulting formula intuitively matches the expected result from considering reactivity to be defined as the fraction of adduct observed at each position in an RNA at the end of a chemical probing experiment. We discuss assumptions and implementation of the model, as well as ways in which the model may be experimentally validated.

## Introduction

Chemical probing has developed into a powerful experimental approach to interrogate RNA structures *in vitro* and *in vivo*^1–46^. In these experiments, chemical reactions between an RNA and a probe creates covalent adducts at positions in the RNA in a pattern that is determined in part by the underlying structure of the RNA^47^. Uncovering the distribution of adduct positions across a population of RNAs is then a means by which to measure structural properties of those RNAs.

Recently, a collection of experimental techniques have been developed that use sequencing technologies to recover the adduct distribution of chemically modified RNAs as accurately as possible in order to infer RNA structures^9, 10, 18, 19, 29–32, 34–37, 40, 43, 44, 46, 48, 49^.

These techniques all use indirect methods to detect adduct positions, since direct detection of chemical probing adducts on an RNA molecule has not been shown to be possible with current sequencing technologies. The most convenient indirect method is to first convert the modified RNA into a DNA molecule using an enzymatic process called reverse transcription (RT). In this process, a reverse transcriptase enzyme (also referred to as RT) catalyzes the synthesis of a complementary DNA (cDNA) molecule in a 3′ → 5′ direction (Fig. 1). When RT encounters an adduct, one of two scenarios is possible: either the RT stops 1 nt before the adduct^50^ (at site *k* − 1) which we call an RT-stop, or the RT proceeds through the adduct and introduces a mutation at site *k*, which we call an RT-mutation.

**Figure 1.**
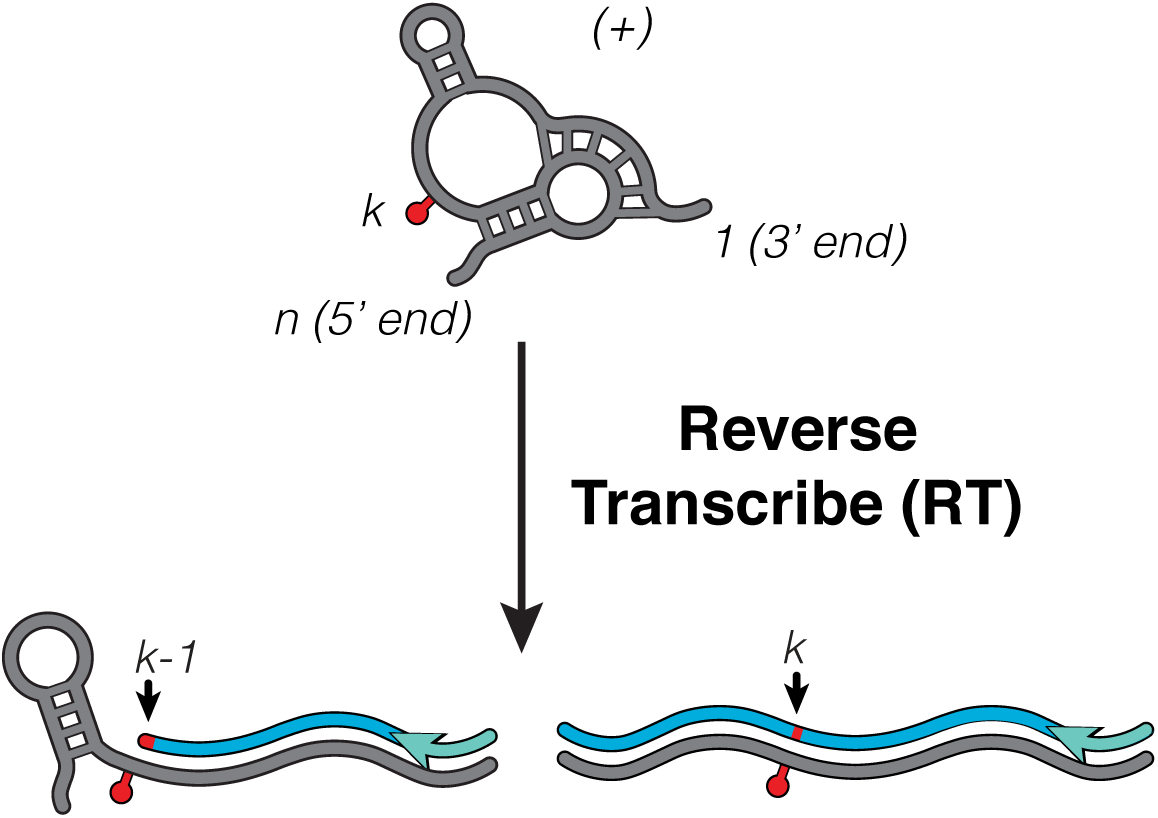
Reverse transcription (RT) encodes the positions of RNA adducts as either stops or mutations. RT converts an RNA (grey) into a complementary DNA (cDNA, blue) in a 3′ → 5′ direction. (A defined RT priming site is shown in teal). If RT encounters an adduct at site *k* (red pin), one of two scenarios is possible: either the RT falls off 1 nt before the adduct (at site *k* – 1, indicated as a red segment) to generate a *k*-fragment (left), or the RT proceeds through the adduct and intruduces a mutation at site *k* (indicated as a red segment, right). (+) refers to these RNAs being present in a population that has already been modified by the chemical probe to form covalent adducts. There are a range of RT priming strategies that can be used with this approach including those that can recover adduct positions at the 3’ end of the molecule^14, 18, 19, 31, 32, 36, 43, 44, 46, 51–57^.

Subsequent steps in the experimental protocols then analyze cDNAs for signatures of RT-stops and RT-mutations. Early versions of the experiments analyzed cDNAs by separation techniques such as capillary electrophoresis that enabled the identification of RT-stops as distribution of different length cDNAs^22, 24, 26, 27, 58–64^. Later innovations developed high-throughput sequencing (HTS) methods to analyze cDNAs^10, 18, 19, 40^. These experiments generally involve a series of ligation, size selection, and PCR steps in order to format cDNAs into a sequencing library with vendor-specific adapter sequences prior to sequencing^65^.

The earliest versions of HTS experiments mapped RT-stops by mapping cDNA ends, and importantly enabled chemical probing to be performed on complex mixtures of RNAs since cDNA signatures from different RNAs could be distinguished through bioinformatic sequence alignment^18, 66, 67^. Later innovations showed that HTS approaches could also be used to map RT-mutation sites through bioinformatic analysis of sequence mutations^32, 68^. These sequencing-based approaches represent an important enhancement in the amount and speed in which adduct detection information could be gleaned from these experimental approaches.

Once adduct signatures are detected and collected as distributions of RT-stops or RT-mutations across each nucleotide of the corresponding RNA sequence, they can then be used to estimate a value for the ’reactivity’ of each nucleotide to the chemical probe. Reactivities contain information about the underlying RNA structure, and specific reactivity values are a result of differences in the propensity of the chemical probe to react with bases that differ in structural context^69, 70^. The main goal of data analysis procedures then is to estimate these underlying reactivity values as accurately as possible given the observed distributions of RT-stops and RT-mutations.

Historically, chemical probing experimental and data analysis approaches used the signatures from RT-stops to estimate reactivities at each position^66, 67^. More recently, groups have begun push to use RT-mutations in order to define reactivity at a given position. These groups argue that inherent enzymatic biases in library prep cause distortions in RT-stop data that lead to calculated reactivities that do not directly correspond to the intrinsic reactivity of a given position. However, this assumes that the information given by RT-stops is correlated with information given by RT-mutations. However, recent papers^52, 71^ have shown not only that these metrics are poorly correlated but also, in some contexts, completely orthogonal. Specifically, recent analysis of DMS probing RT-stop and RT-mutation signatures on the same pool of modified RNAs show that different reverse transcriptase enzymes and reaction conditions show distinct biases for detecting adducts as either RT-stops or RT-mutations^52, 71^. The orthogonality of RT-stops and RT-mutations and the variability between RT-stop and RT-mutations between different enzymes strongly suggest that approaches that only incorporate either RT-stops or RT-mutations miss information about adduct distributions and therefore the resulting reactivities may be incomplete and lacking in accuracy.

Conversely, these observations suggest that improvements in chemical probing accuracy can be achieved by incorporating both RT-stops and RT-mutations in the estimation of reactivities. To address this, here we developed a formalism for estimating chemical probing reactivities using both RT-stops and RT-mutations in a single experiment. Following the work of Aviran *et al.*^66, 67^, we extend a maximum-likelihood derivation of reactivities and present a reactivity formula that uses this combined information. Interestingly, this formula matches an intuitive interpretation of chemical probing reactivities as the fraction of adduct formed at each nucleotide at the end of the probing reaction. We discuss assumptions of this model, and end with a discussion on experimental approaches to validate this model.

## Results

### Model Setup

For an RNA of length *n*, we define the very 3’ nucleotide of the RNA as position 1, and the very 5’ nucleotide of the RNA as position *n*, and only consider cDNAs that start at position 1 (Fig. 1). We define a *k*-fragment as a cDNA whose 5’ end begins at and complements position 1 of the RNA, and whose 3’ end complements position *k* − 1 of the RNA. We emphasize that we **assume that RT-stops and RT-mutations at a given adduct position in a single RNA molecule are mutually exclusive events: since an RT that stops due to an adduct at position** *k* **stops at position** *k* − 1**, it cannot introduce a mutation at position** *k*. This will be an important feature of the formula for recovering reactivities from observations of RT-stop and RT-mutation events, and we discuss implications for this assumption below.

The outcome of a typical high throughput sequencing (HTS) experiment to map RNA chemical probing adducts is a series of cDNA sequences that encode the locations of RT-stops and RT-mutations. These raw cDNA sequencing reads are then processed through read alignment software to generate a list of RT-stops and RT-mutations at each position within the target RNA. To keep track of these observed patterns, we define several variables that indicate stops (*S*) and mutations (*M*).

Since an RT that stops due to an adduct at position *k* stops and transcribes through position *k* − 1, we define 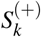 as the number of observed cDNA fragments whose 3’ ends map to position *k* − 1 and 5’ ends map to position 1, where (+) indicates these fragments were observed from samples that had been treated with the chemical probe, called (+) channel samples. We additionally call 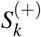 the number of *k*-fragments because information on adduct formation would come from position *k* even though the sequenced length is of *k* − 1. Note that if an RT does not stop internally, then RT will transcribe through the end of the cDNA and reach position *k* = *n*. Thus, 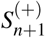 represents the number of full-length reads observed in the (+) channel. Similarly, we define 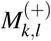 as the number of *k*-fragments observed that have at least one mutation and includes a mutation at position *l* in the (+) channel, where *l < k*.

It is also possible that RT stops or mutates at positions due to natural processes and/or there are cDNA sequencing errors that are not caused by chemical probe adducts. These events confound the measurement of adduct distributions and must be accounted for in data analysis to extract their confounding influence. To do this, control experiments are run that process the RNA in the same manner, but do not include the addition of the chemical probe. Such (-) channel experiments then generate a set of observed RT-stops and RT-mutations which we denote as 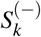 and 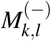, respectively.

Figure 2 shows several examples of possible RT-stop and RT-mutation scenarios in the (+) and (-) channel and how they would each contribute to 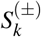 and 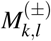.

**Figure 2.**
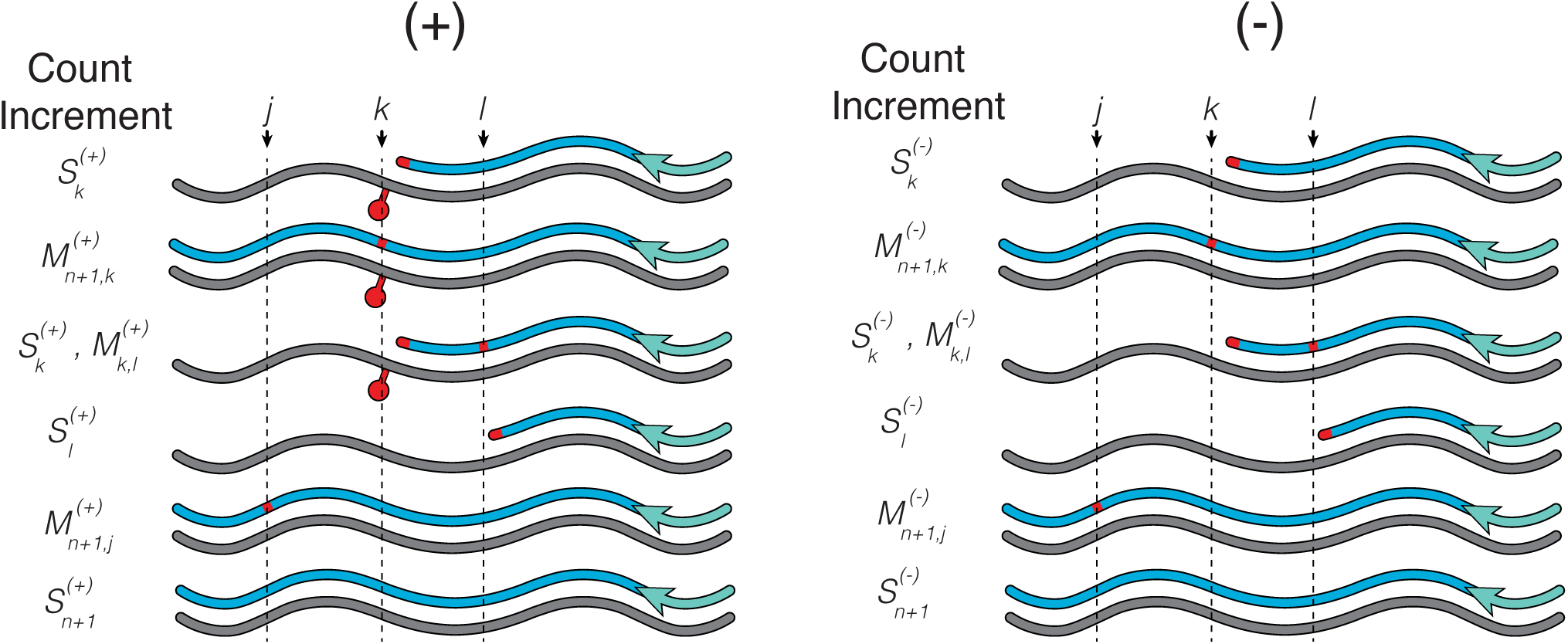
Examples of RT-stops and RT-mutations in the (+) and (-) channels and how they contribute to 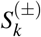 and 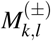. Note that a single read may contribute to both stop and mutation counts. Adduct positions in the (+) channel are denoted by red pins.

Once the raw data is processed, our goal is to use the observed patterns of stops and mutations, 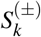 and 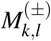, in a formula that more completely and accurately estimates the reactivity of each nucleotide of the interrogated RNAs to the chemical probe. Below we state the results of our derivation of this formula and discuss its implementation and limitations.

### Estimating chemical probing reactivity from RT-stops and RT-mutations

We define the ’reactivity’ of site *k* in an RNA molecule, *r_k_*, as the probability of an adduct forming at that site during a chemical probing experiment. *r_k_* contains structural information about the molecular fold of the RNA sample and is the primary data we want to extract from the probing experiment. Since RT can both stop and mutate at adducts, we also define *β_k_* to be the probability that RT stops due to an adduct at site *k* following syntax in^66^, and *µ_k_* to be the probability that RT mutates due to an adduct at site *k*. Recent evidence suggests that there are strong context preferences for RT to favor either stops or mutations at a given position of an RNA, and so both must be accounted for in our estimate of *r_k_*^52, 72^. Since RT-stops and RT-mutations are mutually exclusive events when detecting adducts, we define the reactivity at site *k* to be the sum of these two probabilities: *r_k_* = *β_k_* + *µ_k_*.

Below we present a maximum-likelihood framework derivation following^66^ for generating our best estimate of these probabilities 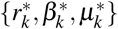 given the observations of 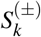 and 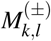. Using this framework we obtain

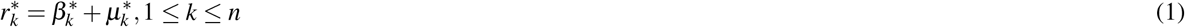

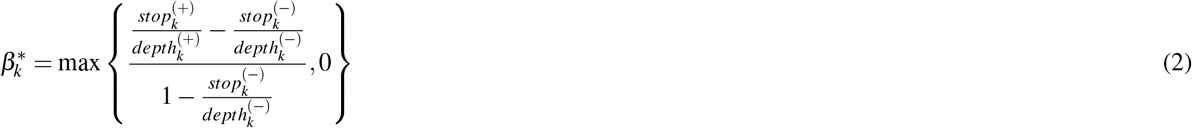

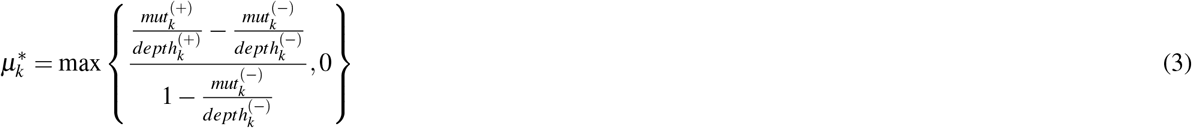

where 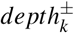 is the number of sequencing reads that cover position *k* in each channel, which can be calculated from

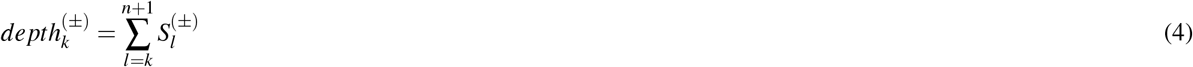

and the mutations observed at position *k*, *mut_k_* is defined as

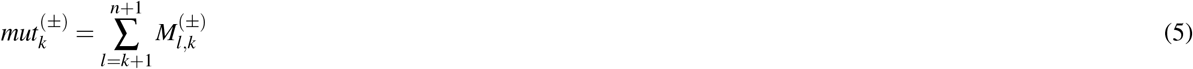

We rewrite 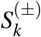 as 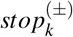 in Eq. 2 to more clearly delineate RT-stop and RT-mutation information.

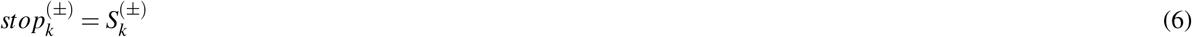

The quantities 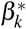 and 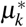 are the best feasible estimates for the reactivity components due to RT-stops and RT-mutations, respectively. As explained below, they come from 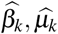 which are the estimated parameters from observed data. Since 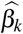 and 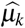 are each calculated as the difference between terms calculated from the (+) and (−) channel data, there is no guarantee that they will be strictly nonnegative. Therefore, in the case that the calculation results in 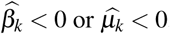, the feasible solution is to set them to zero^66^, hence the use of the maximum function in defining 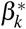 and 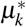. Note this arises in cases when there is severe dropoff or mutations in the (−) channel indicating a large amount of noise in the measurement at this position. It is therefore adviseable to keep track of scenarios where 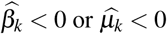 since they are generally indications of poor data quality, especially for larger negative values, where experimental conditions may potentially be optimized.

## Discussion

### An intuitive link between reactivity estimates and fraction of adduct formed

Interestingly Eq.1 has an intuitive interpretation that is related to the concept of the fraction of adduct formed at the end of a chemical probing reaction. Since *depth_k_* is an indication of how many adduct detection events are possible at each position, both RT-stops and RT-mutation data are incorporated in terms that have the form

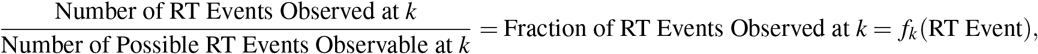

where RT Event refers to either RT-stops or RT-mutations. Therefore we can write

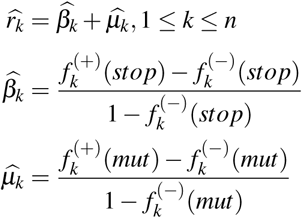

Thus both 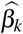 and 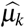 represent the fraction of RT events observed in the (+) channel corrected for the fraction of RT events observed in the (-) channel for stops and mutations, respectively. The denominators in each term represent the fraction of signal due to adduct that is possible to observe in the (+) channel. These denominators arise because an RT event observation can be due to an adduct *or* a background process, but not both^66^ – i.e. if a fraction of RT events is observed in the (-) channel, the fraction of events that then *can* be observed in the (+) channel is reduced by that amount in order to estimate the fraction of events due to true signal. The denominators effectively correct for scenarios in which there is high background that obfuscates signal due to adducts. In cases where there is little or no background, these denominators can be approximated to be ∼ 1 and we have

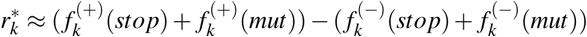

Since the (+) channel has signal due to adducts and background processes, while the (-) channel only has signal due to background processes, the subtraction amounts to

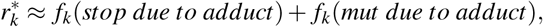

where *f_k_*(*event due to adduct*) denotes RT-stops or RT-mutations at site *k* due to adduct and not due to natural fall off and mutations.

Since RT-stops and RT-mutations are the two ways to detect adducts, then

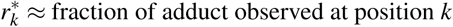

Importantly, the fraction of adduct formed at any given position is a quantity that is determined by the chemical kinetics of the probing reaction and the structure-dependent fluctuations of each nucleotide of the RNA^47, 69^. By estimating reactivities that correspond to the fraction of adduct observed, reactivity values should most closely align with the kinetics of the chemical probing reaction, which should allow a deeper understanding of data from high throughput RNA structure probing experiments.

### Model Assumptions and Implementation

Two main assumptions were made in the above model:

1. Mutations at different positions are independent of each other.
2. Observing an RT-mutation at a given position is exclusive to observing an RT-stop at that position.

The first assumption is reasonable for low modification rates. However, some studies^47, 73^ indicate that the chemical adducts caused by certain probe reagents may alter the structural dynamics of the RNA once formed, which could in turn influence the formation of additional adducts if high modification rates are used - i.e. that certain chemical probes could potentially destabilize individual RNA molecules such that additional adduct formation to the same molecule would not reflect the native structures of interest. More work is needed to understand how multiple probe modifications can impact the ability to estimate reactivities which may depend on the nature of the probe, where it chemically modifies the RNA base, and any sequence/structural contexts of these scenarios.

The second assumption is more nuanced and impacts the implementation of the read mapping and application of the above reactivity formulas. In particular, the case of the very 3’ end of a cDNA presents a challenging mapping case if it is mutated. According to the assumption, a mutated cDNA 3’ end at position *k* − 1 would contribute to two counts: *S_k_ and M_k,k_*_−1_. However, it could be possible that in the process of stopping, RT introduces the mutation at the same time and thus this event should only be counted once. More work on biochemically defined adducts would be needed to validate or modify this assumption.

### Experimental Validation

The major innovation in the above formula for chemical reactivities is to formally incorporate the observed signatures of *both* RT-stops and RT-mutations when estimating reactivities. The major motivation for this is the data and results of^52, 72^ which clearly show that RT drop and mutation signatures on the same pool of modified RNA differs depending on the specific RT enzyme and associated buffer conditions used to perform the conversion into cDNA. The goal of the RT process to convert every RNA adduct into a detectable signature in cDNAs therefore justifies our assertion that both RT-stops and RT-mutations should be included simultaneously in the reactivity estimation.

While the above formula for chemical probing reactivities makes intuitive sense as the fraction of adduct detected at each position, it still requires experimental validation. Accordingly, we expect two important improvements from applying the combined RT-stop+map model: improvement in chemical probing reactivity accuracy, and an invariance of reactivities to RT conditions.

Improvements in chemical probing reactivity accuracy are naturally expected since current approaches that focus solely on RT-stops^3, 11, 18, 30, 34, 36^ or RT-mutations^32, 46^ will inherently miss information^52, 72^. Many approaches to assess chemical probing accuracy rely on an indirect method to first utilize reactivities in RNA secondary structure prediction algorithms, and then assess accuracy of reactivity data based on the improvements in structural prediction^74–77^. While we expect that RT-stop+map reactivities may improve the accuracy according to this benchmark, we anticipate that improvements may only be modest as it appears that the current RNA structure prediction algorithms are starting to reach the inherent limits of their accuracy given the assumptions used in their calculations^78^. Therefore methods that assess accuracy of reactivities through more direct analysis and/or new RNA structure prediction algorithms may be needed to uncover improvements when using both RT-stops and RT-mutations.

The other improvement suggested by the RT-stop+map model is an invariance of reactivities to RT conditions. In other words, estimated reactivity values should be the same no matter what RT enzyme or RT conditions are used. This is because the RT process is a means to detect RNA adducts. If the same pool of modified RNA is used, then this adduct distribution will not change, making it a goal of adduct detection methods to uncover the same distribution independent of the method conditions. Interestingly, this invariance is also suggested when observing the individual RT-stop and RT-mutate reactivities from Sexton *et al.*^52^ and Novoa *et al.*^71^, which strongly suggest that adding the two together would create reactivities that are highly similar between RT enzymes and conditions.

While invariance to RT conditions is strongly suggested as an outcome when using both RT-stops and RT-mutations, it is not guaranteed, mainly because it should only be true when reactivity estimates converges to the true fraction of adduct formed value. This can breakdown for simple reasons such as inadequate sequencing depth needed to overcome high background stops and mutations, or more complex reasons related to biases in specific library preparation steps. In particular, biases introduced by ligation or PCR steps that prevent adducts in specific sequence contexts to be uniformly sampled would interfere with more accurately estimating reactivities. More work is needed to test different experimental library preparation protocols in the context of the RT-stop+map reactivity estimation in order to examine these effects. Interestingly, searching for library preparation strategies that are invariant to RT conditions may be a means to identify the most accurate experimental strategy. Thus, future work in this area may produce useful insights in both experimental protocols as well as more accurately estimating reactivities.

## Conclusion

In this work, we derive an equation for estimating chemical probing reactivities that uses information from both RT-stops and RT-mutations. This is based off of recent work^52, 72^ that gives strong evidence that RT-stop and RT-mutation detection methods give complementary information when used with DMS probing - i.e. each method has context dependence such that they tend to map adducts in unique scenarios rather than mapping the same adduct positions. Therefore, we propose that reactivity estimation that considers both RT-stop and RT-mutation will be more accurate than methods that consider only one source of adduct detection. Future work will require the above formulas to be tested in a range of experimental contexts to demonstrate that the conclusions drawn from it are robust. We hope that these efforts will lead to improvements in RNA structure interrogation methods that are becoming increasingly important in answering questions about how RNA structure impacts a broad range of processes in biology.

## Full Derivation

### Model Setup

Chemical probing reactivities represent probabilities that adduct formation will occur at each nucleotide in an RNA during the probing reaction. Once the reaction proceeds to completion, these probabilities will naturally manifest themselves as a fraction of adduct formed at each position, which is defined as the proportion of those nucleotides that have the adduct out of the total population.

A given pattern of reactivities will naturally generate a distribution of RT-stops and RT-mutations across a population of cDNAs when the RNAs are reversed transcribed. Thus given a set of known reactivities, an observed pattern of RT-stops and RT-mutations could be calculated. However, in chemical probing experiments the information available is the converse - we know the pattern of RT-stops and RT-mutations, but do not know the underlying reactivities that gave rise to those patterns. The process of deriving a formula to estimate reactivities is thus to search over all possible reactivity values that lead to RT-stop and RT-mutate distributions that most accurately match the observed data. Fortunately this can be done exactly to yield a closed-form expression for the most accurate reactivity values possible from observed patterns of RT-stops and RT-mutations.

The derivation of equation (1) follows the maximum likelihood approach originally developed in^66^ for the case of just detecting RT-stops. The maximum likelihood approach describes adduct detection by RT as a probabalistic process, where as RT processes through cDNA synthesis there are certain probabilities for it to fall off or mutate due to either encountering an adduct or due to natural processes. We define the probabilities

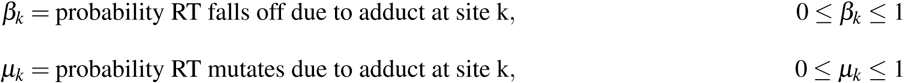

where the ranges for *β_k_* and *µ_k_* are set since they are probabilities. When describing a complete model, we also need to account for the probabilities for RT to fall off or mutate due to natural processes, which are described by two additional probabilities

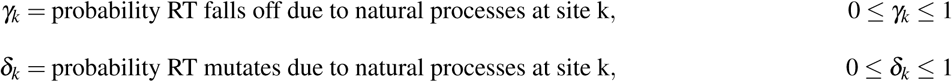

where again ranges for *γ_k_* and *δ_k_* are set since they are probabilities. If known, these probabilities define the reactivity information we desire from the chemical probing experiment from equation (1). However, for a given RNA these probabilities are unknown, and it is the goal of the maximum likelihood framework to estimate these probabilities given the information obtained from the sequencing reads. Estimated parameters from the maximum likelihood estimation are then 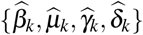 which we then enforce nonnegativity to obtain the estimated probabilities 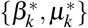 and thus the estimated reactivity 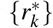. For shorthand we define *B* = {*β_k_*}, Γ = {*γ_k_*}, *M* = {*µ_k_*}, ∆ = {*δ_k_*}.

### Construction of the Likelihood Function

Even though the probabilities {*B, M,* Γ, ∆} are initially unknown, we can still use them to construct the overall probability of observing a specific type of sequencing read in the experiment. For example, if we only consider RT-stops, the probability that we would observe a *k*-fragment in the (-) channel would be

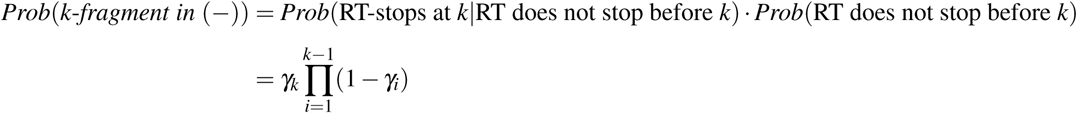

The first term in the equation is the probability RT stops at position *k*, which is *γ_k_*. The second term is the probability that RT does *not* stop before reaching position *k*. Since the probability of *not* stopping at a position *i* is (1 − *γ_i_*), then the probability of not stopping before *k* is the product of all the (1 − *γ_i_*) terms where *i < k*. Note that since we were describing events in the (-) channel, we use Γ = (*γ*_1_*, …, γ_n_*) since they describe RT stop events due to natural processes which are the only causes of RT stop events in the (-) channel. Following this logic, we can write the probability of observing a full length fragment in the (-) channel as the product of RT not stopping the whole length of the molecule, or

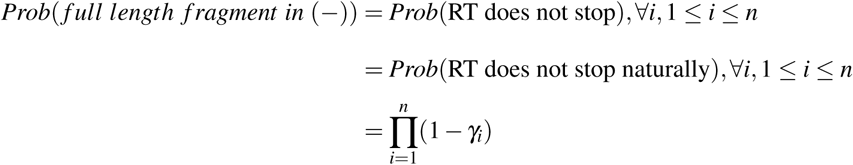

When considering RT-stops in the (+) channel, things are slightly more complex since RT can stop both due to adducts present *and* natural processes as well. Thus when we write down probabilities for observing fragments in the (+) channel, these probabilities will involve both *B* = (*β*_1_*, …, β_n_*) and Γ = (*γ*_1_*, …, γ_n_*). When considering the probability for observing a full length fragment in the (+) channel, we simply need to include (1 − *β_i_*) with (1 − *γ_i_*) in the probability for RT *not* stopping at position *i* leading to

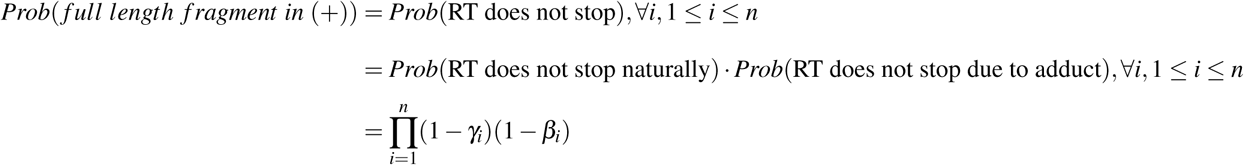

For the probability of observing *k*-fragments in the (+) channel there are two independent causes for dropoff at *k*: either the *k*-fragment was due to an adduct at *k* or due to a natural dropoff at *k*:

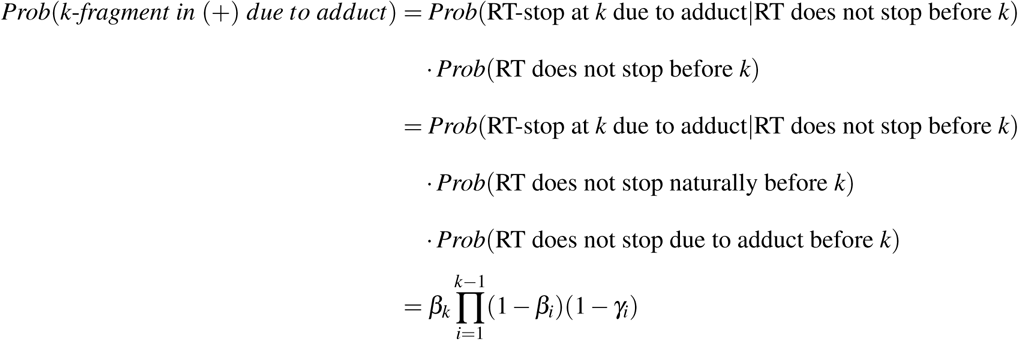

Here, the first term in the equation is the probability RT stops at position *k* due to adduct, which is *β_k_*. The second term is the probability that RT does *not* stop before reaching position *k*, and since the probability of *not* stopping at a position *i* in the (+) channel is (1 − *β_i_*)(1 − *γ_i_*) (which is the probability of not stopping due to adduct *and* not stopping due to natural processes), then the probability of not stopping before *k* is the product of all the (1 − *β_i_*)(1 − *γ_i_*) terms where *i < k*.

RT can also fall off in the (+) channel due to natural processes, so we must account for this probability as well:

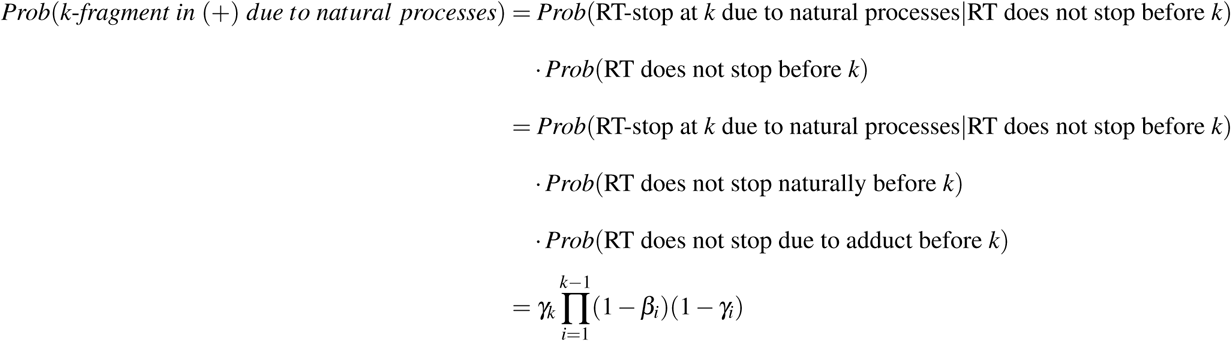

The first term, *γ_k_* represents the probability of falling off due to natural processes. The other term in this equation describing the probability of no RT-stop before *k* remains the same.

However, we must disentangle RT fall off in the (+) channel due to natural processes versus adduct formation. This is particularly important for positions that have both a tendency for natural process RT-stops as well as adduct formation. In other words, a position *k* with the probability of adduct formation *β_k_ <* 1 could also create a *k*-fragment from natural processes as governed by the probability *γ_k_*. We account for this in the following equation:

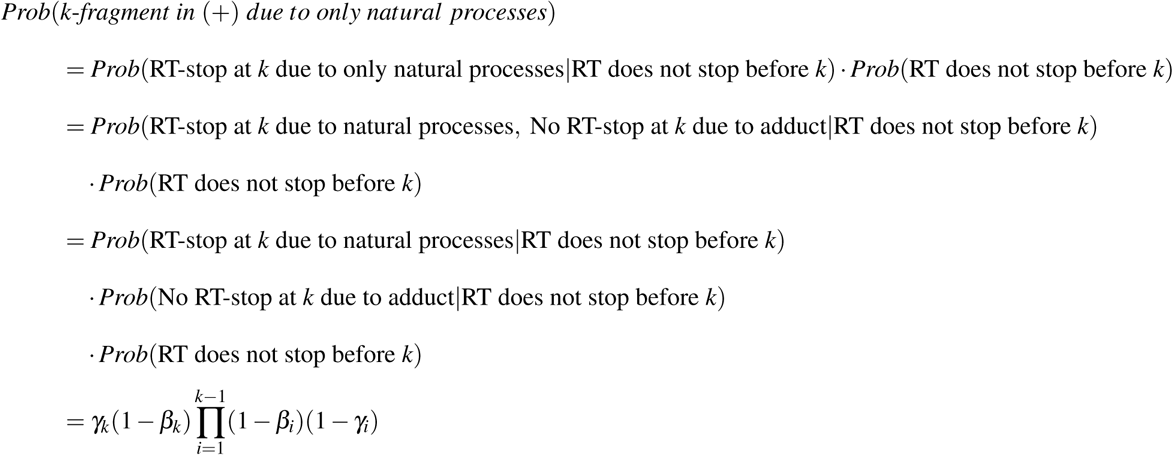

Note the slightly different form of this probability. The first two terms represent the probability of falling off due to natural processes *and not* due to adduct, which is *γ_k_*(1 − *β_k_*). The last term in this equation describing the probability of no RT-stop before *k* remains the same. Since we have taken into account the possible scenarios, to find the probability of a *k*-fragment in the (+) channel due to either type of process, we simply sum them:

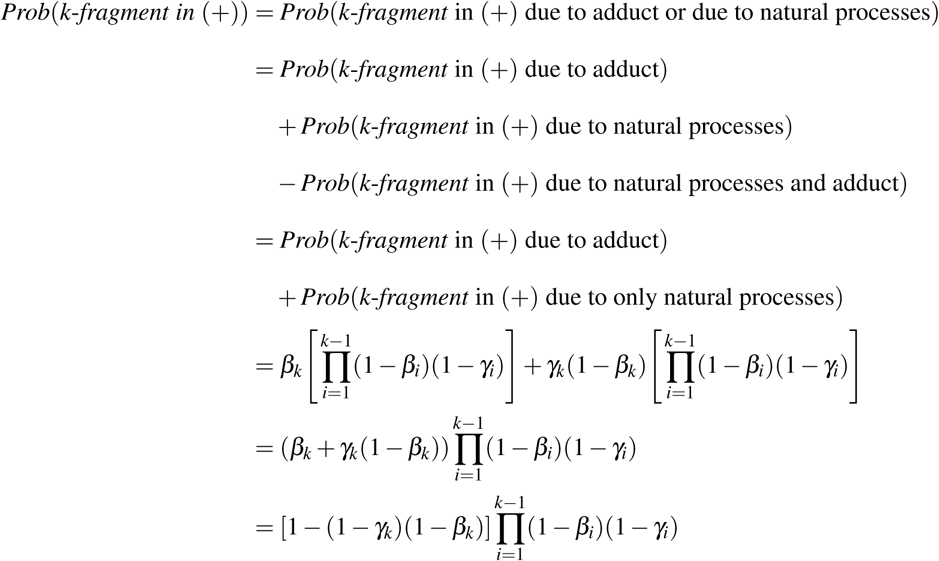

The first term in square brackets in this equation represents the nature of RT falling off due to an adduct or a natural processes. Here (1 − *γ_k_*)(1 − *β_k_*) is the probability of not falling off due to a natural process *and* not falling off due to adduct. Therefore 1 − (1 − *γ_k_*)(1 − *β_k_*) is the probability of falling off due to one or the other. The last term in this equation describing the probability of no RT-stop before *k* remains the same.

The RT-stop-only terms are the same as used in^66^ to derive a reactivity formula in terms of RT-stop events. Here we extend this to include the observation of mutations in cDNA products as well, which are governed by the {*M,* ∆} probabilities. Therefore we must extend the probability terms above to account for the different scenarios of observing specific patterns of mutations across the cDNA molecules. There are two important aspects of RT-mutations that we need to incorporate when constructing these probabilities. The first is the *assumption* that an RT-mutation at position *k* is *mutually exclusive* to an RT-stop at the same position. This is reasonable because RT stops one nucleotide *before* the roadblock it encounters (either adduct or some interfering element that contributes to natural drop off). Therefore an RT-stop due to a road block at position *k* results in a cDNA that ends at position *k* − 1, which could therefore not have a mutation at position *k*. This mutually exclusive nature modifies the probabilities for observing a *k*-fragment in the (-) channels in the following way:

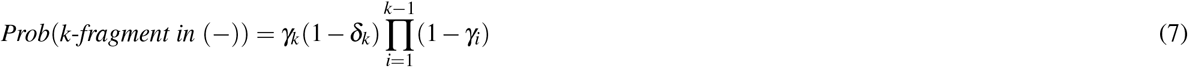

where the *γ_k_*(1 − *δ_k_*) term reflects the mutually exclusive nature that if an RT-stops due to a natural process at position *k* (with probability *γ_k_*), then it cannot introduce a mutation at position *k* (with the probability of not mutating (1 − *δ_k_*)). Similarly, the probability for observing a *k*-fragment in the (+) channel is modified to:

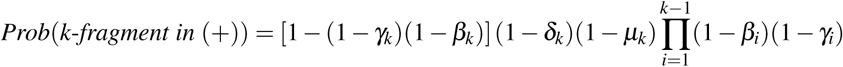

where we have incorporated the probability that there was no mutation at position *k* due to natural processes (1 − *δ_k_*) and no mututation at position *k* due to adduct (1 − *µ_k_*) with the (1 − *δ_k_*)(1 − *µ_k_*) term.

The second aspect of mutations that we need to incorporate is the specific pattern of mutations that may be present across the rest of the cDNA molecule. We *assume* that mutations at different positions are independent of each other – i.e. if a mutation can occur at position *i* due to natural processes with probability *δ_i_*, then the probability of observing mutations at positions *i* and *j* is just the product *δ_i_δ _j_*. If a full length cDNA in the (-) channel only had mutations at *i* and *j* and nowhere else, then the probability of observing this fragment would be 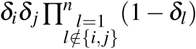 since every other position other than *i* and *j* would not be mutated. In the same way, the probability of observing any pattern of mutations across a cDNA of length *k* − 1 in the (-) channel can be written as:

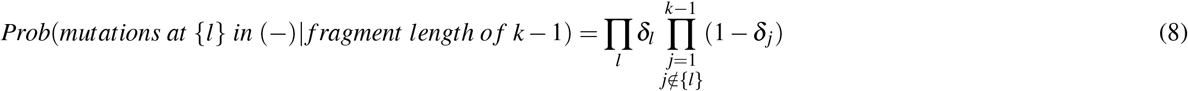

where the notation *j* ∉ {*l*} indicates that the second product covers the positions *j* that are not mutated. For (+) channel cDNAs, we have a similar scenario as with RT-stops – RT-mutate events due to an adduct or natural processes are independent. An RT-mutate event due to natural processes at position *k* occurs with probability *δ_k_*, while that due to an adduct alone occurs with probability *µ_k_*(1 − *δ_k_*). Summing these two then gives the probability that an RT-mutate event occurs at postion *k* in the (+) channel is (1 − (1 − *δ_k_*)(1 − *µ_k_*)).^1^ With this we can write

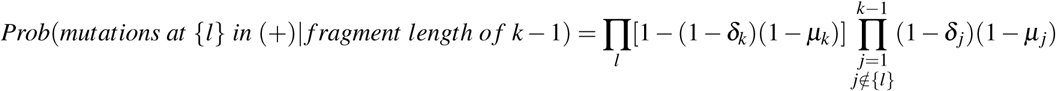

While we have written down all of the different probabilities for observing different types of cDNA fragments, we still do not know the true underlying {*B, M,* Γ, ∆} probabilities. However, we can estimate these numbers given the observed cDNA reads using maximum likelihood estimation. The overall concept of maximum likelihood estimation is that if we can find the set of underlying probabilities 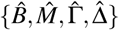 that is most consistent with our observed data, then this will be our best estimate of these parameters. To do so we first construct a likelihood function, *ℒ* ({*B, M,* Γ, ∆}), which represents the likelihood that we would observe a given set of cDNA reads given the set of probabilities {*B, M,* Γ, ∆}. To then find the set of 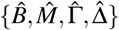 most consistent with our data, we then maximize *ℒ* ({*B, M*, Γ, ∆}) with respect to these parameters given our observed data.^2^

To construct *ℒ* ({*B, M,* Γ, ∆}), we raise the probability of a particular observed event (*p*) to the *N*^th^ power, or *p^N^*, where *N* is the number of times the event was observed. Since we have already constructed the probabilities for observing the different types of cDNA events, we can simply raise these probabilities to powers equal to the number of those types of reads observed. For example from equation (7), we can write

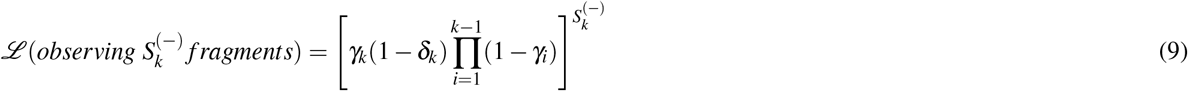

Note however that this does not consider the pattern of mutations that may be present in specific (-) channel *k*-fragments. Since mutations away from the stop site are assumed to occur independently, we can simply multiply the term above to the likelihood of observing a specific pattern of mutations. To construct the likelihood functions for mutations in a *k*-fragment, we must take into account two different observations: the number of *k*-fragments observed with a mutation at a specific position *l*, *M_k,l_*, and the number of fragments observed that were *unmutated* at position *l*, *U_k,l_*. (This is similar to accounting for the number of heads in a coin flip and the number of not heads (tails) in the footnote example.) The latter must be taken into account to account for all observed fragments. Since mutation events are independent, the likelihood of observing *M_k,l_,U_k,l_* is then according to the equation (8)

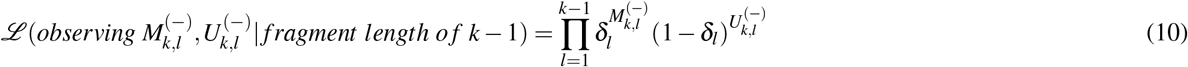

Note the above equation is essentially obtained by multiplying versions of equation (8) together for every cDNA observed. Rearranging all of the different products together will naturally collapse the terms into the form above which raises the probabilities of observing a mutation or not observing a mutation to the number of times those events were observed. Combining equations (9) and (10) we have

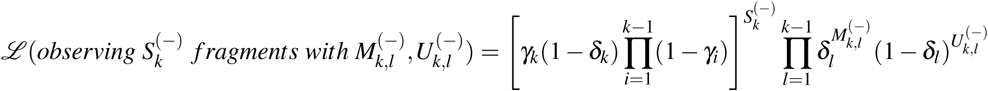

Similarly, for full length fragments in the (-) channel, we have:

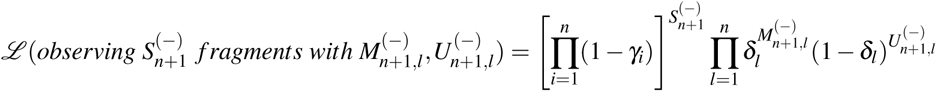

Note the (1 − *δ_n_*) term is incorporated into the right hand side of this equation. The full likelihood function takes into account all of the observed 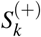 and 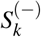 fragments from *k* = 1*, …, n*, as well as the full length fragments 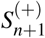. We can therefore write

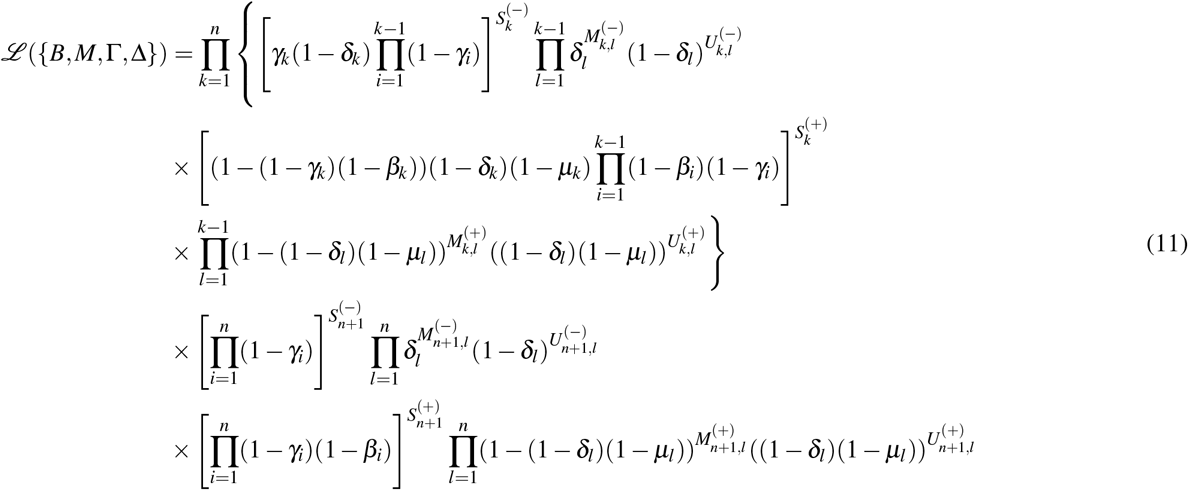

where we have combined terms for *k*-fragments observed in the (-) and (+) channels, and complete fragments observed in the (-) and (+) channels in order.

### Maximizing the Likelihood Function

Our next goal is to maximize *ℒ* with respect to {*B, M,* Γ, ∆} (Eq. 11), given our observations of 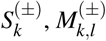, and 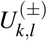. To do this, we first take the logarithm of *ℒ* to separate variables in order to more easily find the ML estimates and obtain:

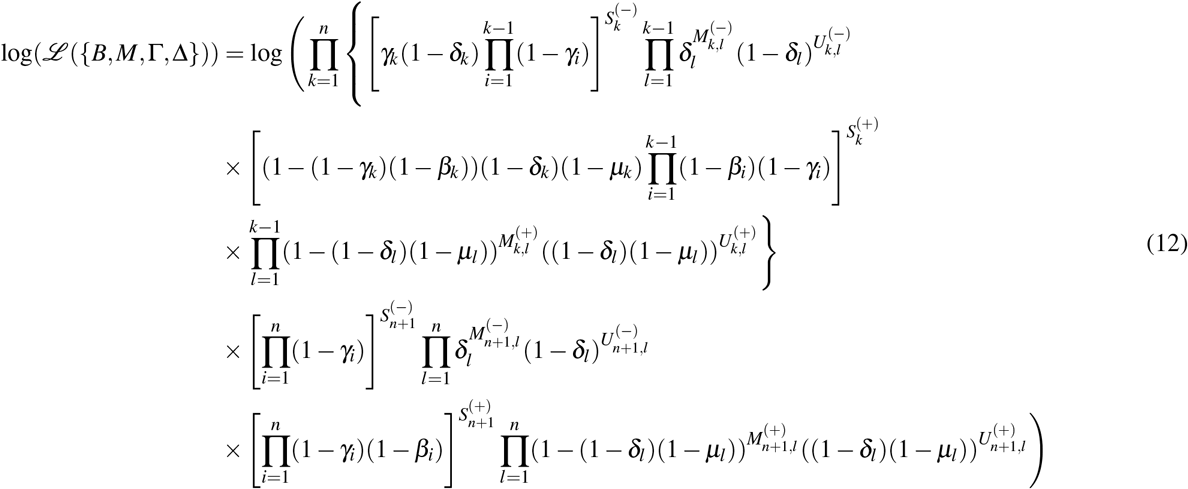

We reduce Eq. 12 further:

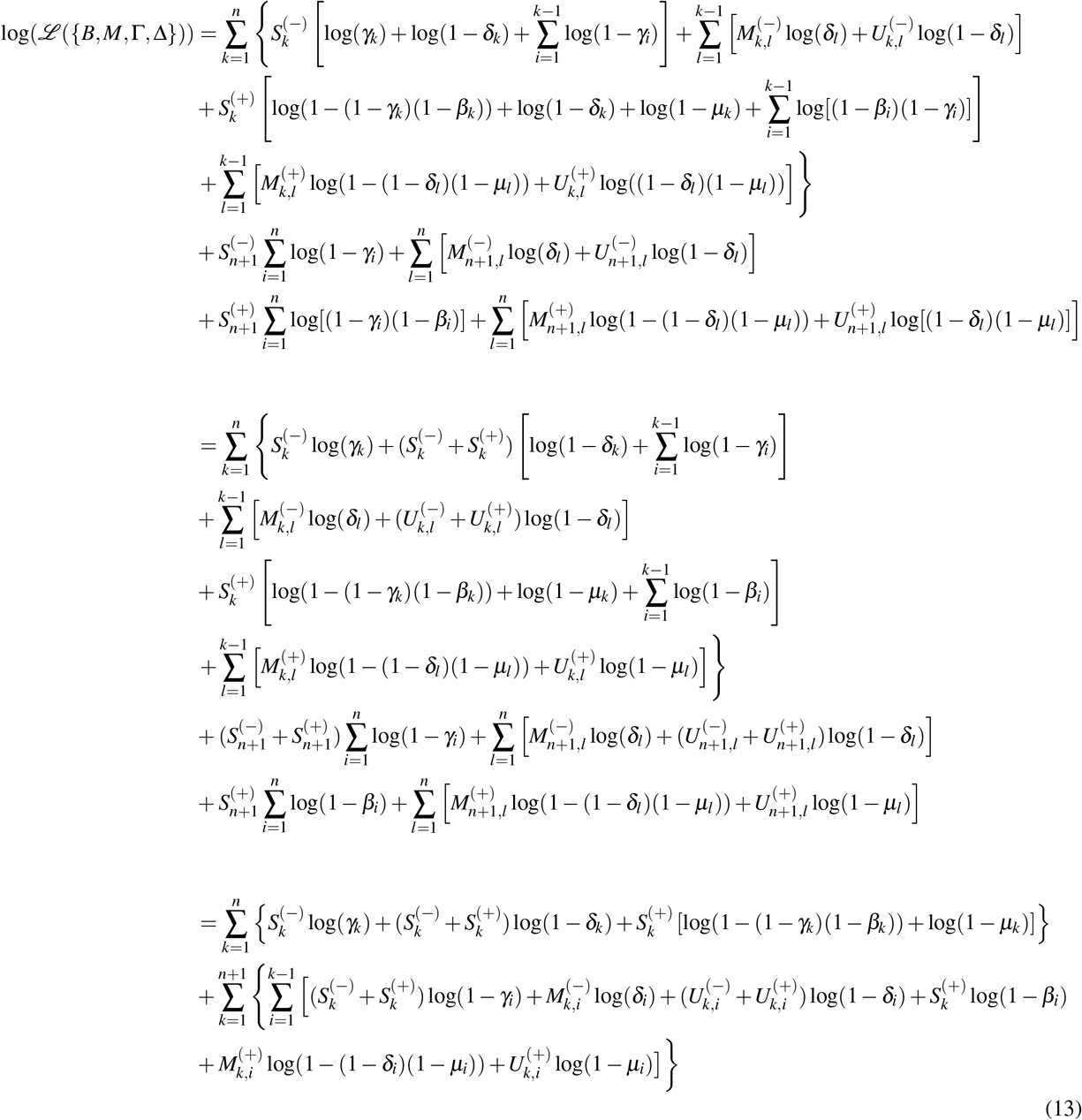

Using the identities 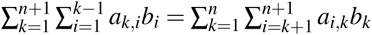 and 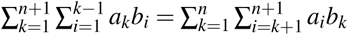 we can rearrange (13) to more easily find the maximum of the likelihood function.

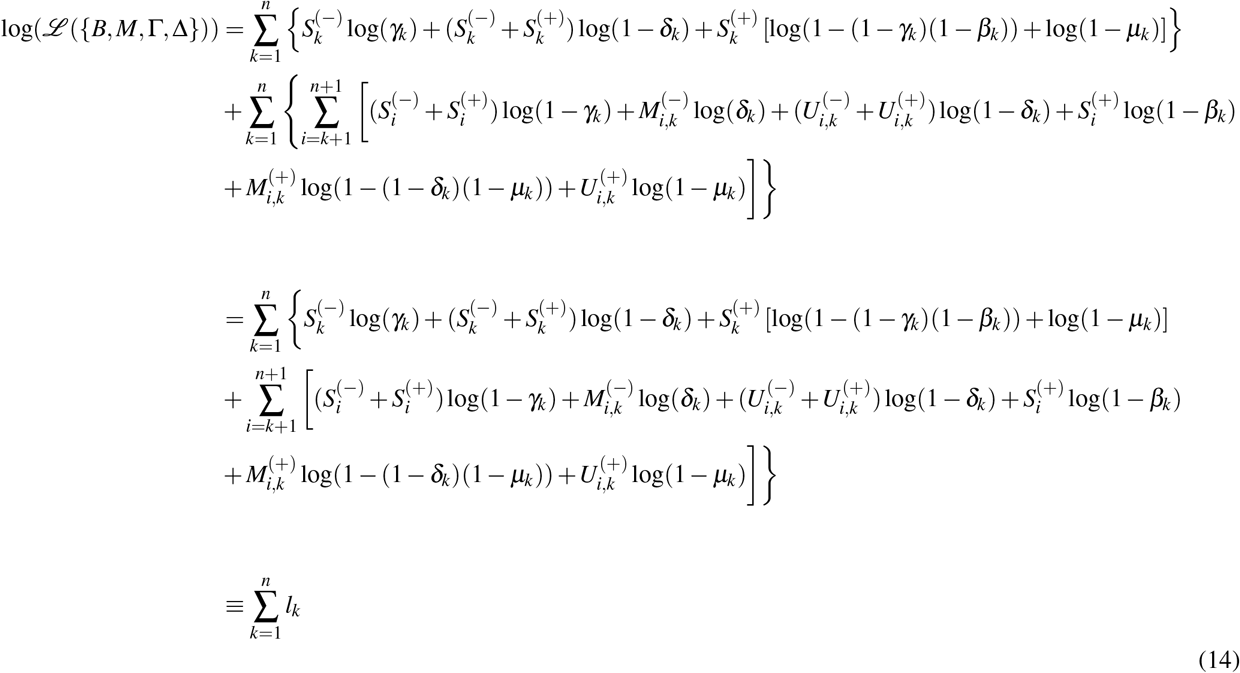

Let 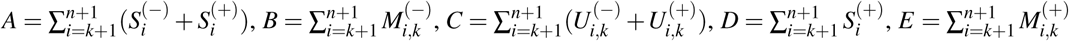 and 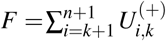. Then from (14):

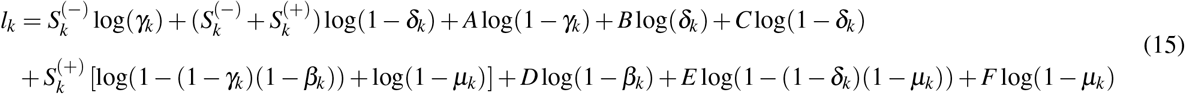

From (15) we can calculate partial derivatives with respect to each parameter in the likelihood function. By setting these paritial derivatives to 0, we can find a critical point for each parameter, which we later show these critical points are maxima and thus are maximum likelihood estimators. Since (14) is a sum of independent terms, we can maximize these independently.

Starting with 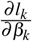:

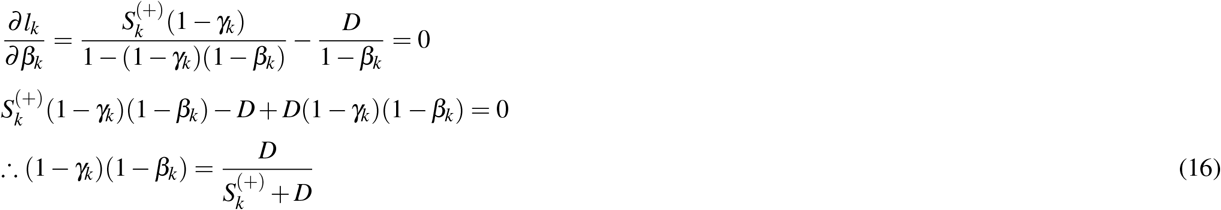

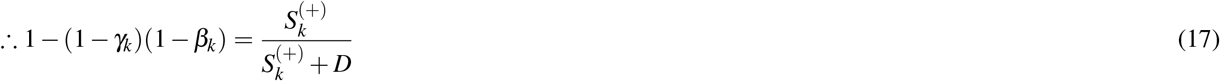

We then use (16) and (17) to calculate 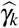 from 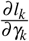.

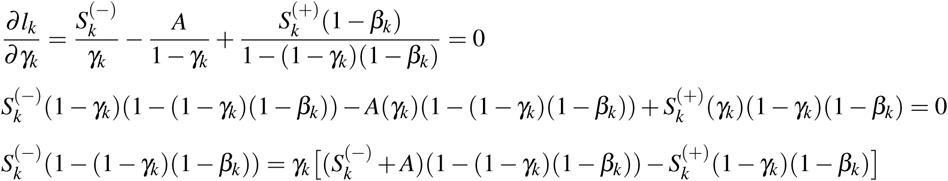

The solution 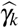 then is

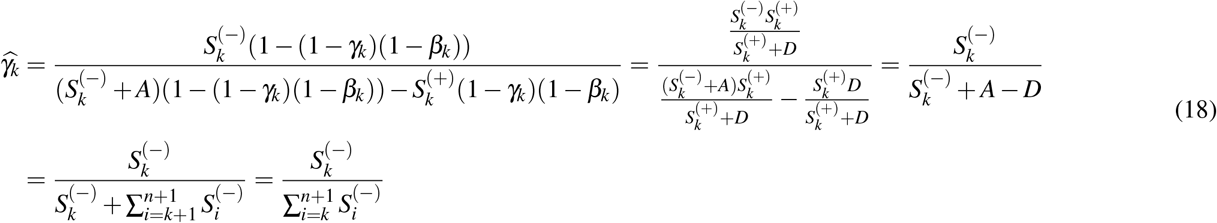

Let 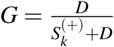. Using *G*, (16), (17), and (18) we can solve for 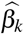:

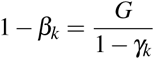

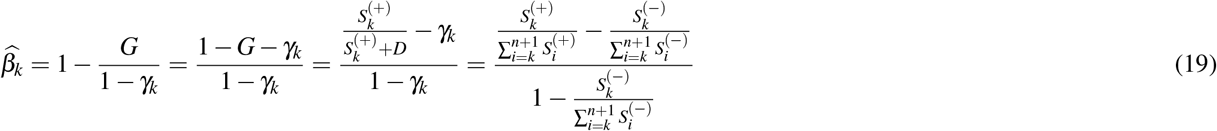

We can solve for 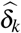 and 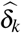 similarly:

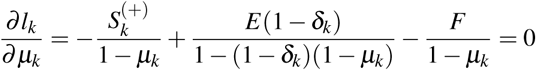

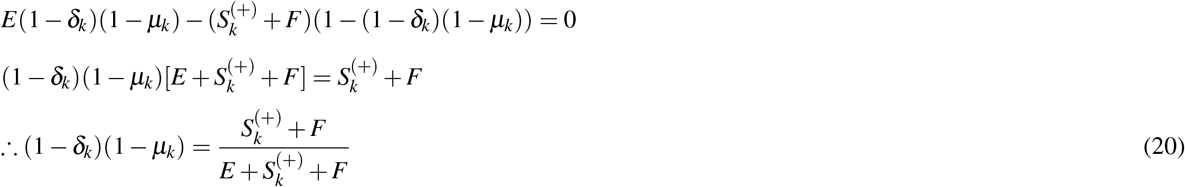

We then use (20) and (21) to calculate 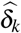 from 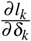.

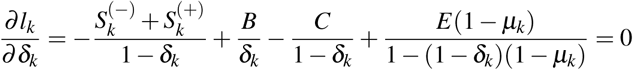

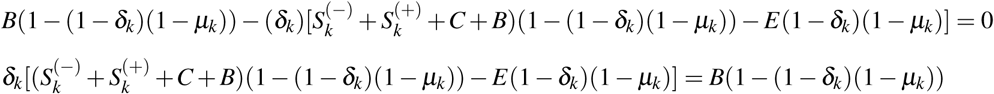

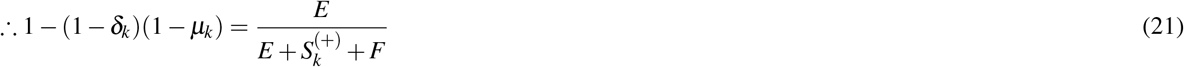

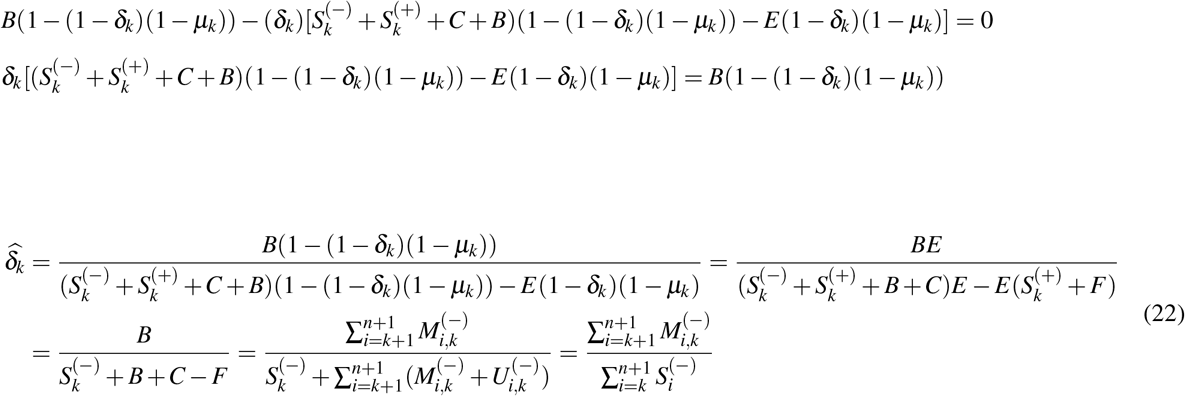

Where we have used 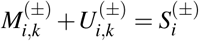 since the former is all *i*-fragments that are either mutated at *k* or not, which is just 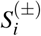. Let 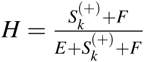. Using *H*, (20), (21), and (22) we can solve for 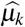:

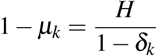

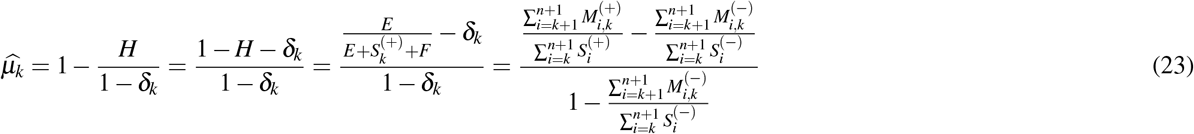

Thus, we have calculated the maximum likelihood estimators 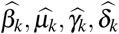 as defined in equations 19, 23, 18, 22.

We additionally use Eq. 5, 4, 6 and propose that the estimated reactivity 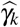 is then:

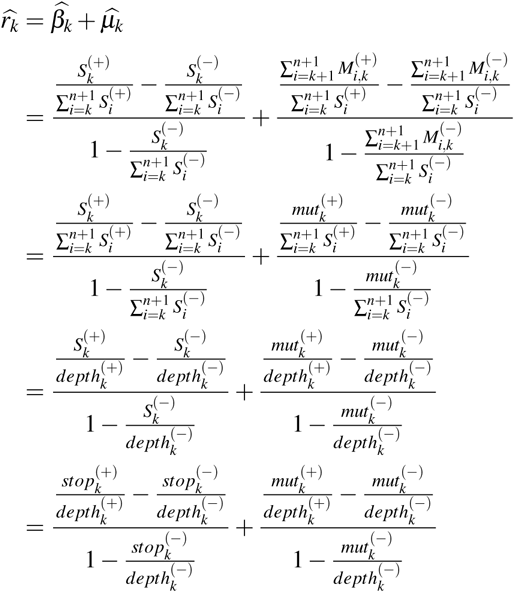

However in practice with real data, 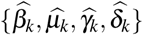 are not guaranteed to be between 0 and 1 inclusive for all *k*. Thus we enforce nonnegativity and our final reactivity calculation 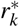 is as outlined above in Eq. 1, 2, 3 and below:

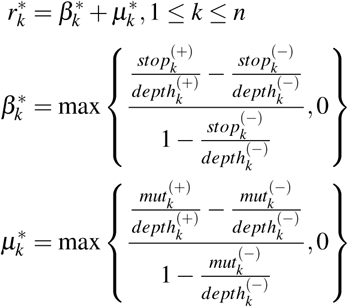

We note the formula for 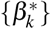 is identical to that of^66^.

### Maximum Likelihood Estimators are Maxima

Next, we show that 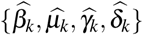 are maxima by taking second partial derivatives of the likelihood equation *l_k_*.

For 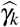:

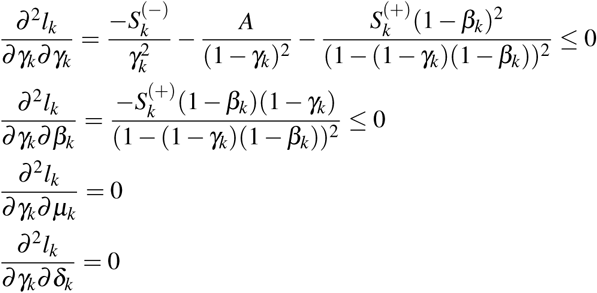

For 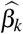

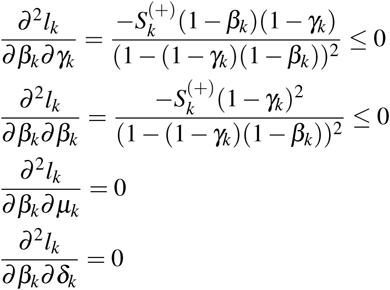

For 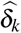

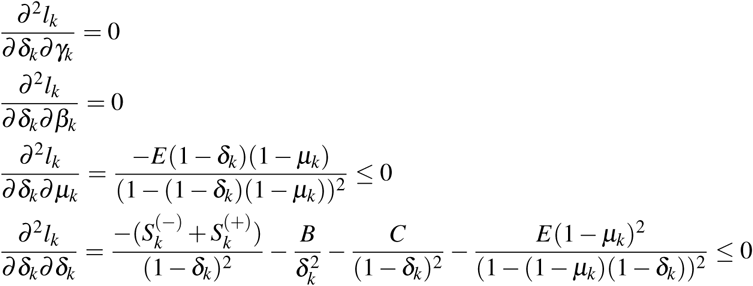

For 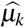

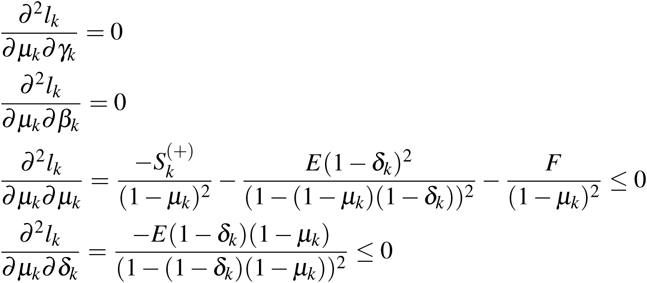

When 0 < β_k_, µ_k_, γ_k_, δ_k_ *<* 1 and 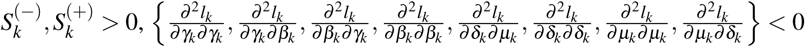 and thus 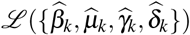 maxima in respect to the variables of these partial derivatives. Otherwise the second partial derivatives are all ≤ 0.

### Estimating the Rate of SHAPE Adduction Formation

Most experimental designs when using RT-stops aim for a single SHAPE adduct formation per RNA present^79^. However, these experimental considerations do not guarantee single hit kinetics so it is useful to estimate the rate of SHAPE adduct formation from the sequencing reads. This estimation then informs the effectiveness of the probing step and its effects on picking up signal. Following^66^ if we assume the probability of SHAPE adduct formation follows a Poisson distribution with rate *c*, then:

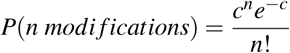

The probability of no modification would then be the following, considering our model both considers RT-stops and RT-mutations:

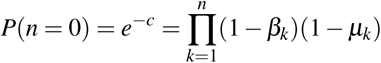

Therefore:

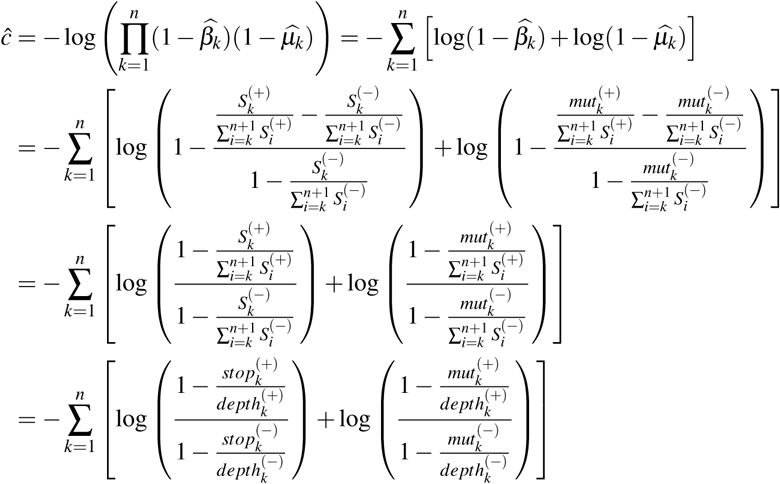

## Acknowledgements

We would like to thank Aaron Coraor for informative discussions about the chemical kinetic view of reactivities, as well as Adam Silverman and Eric Strobel for similar discussions and comments on the detailed derivation. We also thank Chaitan Khosla for inspiring the connections between the chemical and statistical perspectives of reactivities.

## Footnotes

1 Note an easy interpretation of this term is that the probability of not mutating due to a natural process and not mutating due to an adduct at *k* is (1 – *δ_k_*)(1 – *µ_k_*), therefore the probability of mutating at this position is just 1 minus this.

2 The maximum likelihood approach can be explained with the example of a coin flip experiment. Suppose we have a coin that has the probability of *h* for observing a heads and 1 – *h* for observing a tails. Given *h*, the likelihood of us observing *m* heads and *n* tails in a series of *m* + *n* flips is: *ℒ*(*h*) = *h^m^*(1 – *h*)^*n*^. Suppose we do not know *h*, but we have observed *m* and *n*. The question is, what is our best estimate of *h*? We can achieve this by maximizing *ℒ*(*h*), by maximizing log(*ℒ* (*h*)). Since log 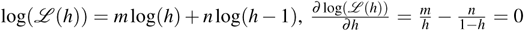, which has the solution 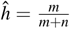. It is easy to show that 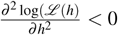 so this is a maximum. Thus our best estimate of *h* is just the fraction of heads observed. The maximum likelihood approach used here to estimate reactivities is just a more elaborate example of this coin flip experiment.

